# Disassembly of embryonic keratin filaments promotes pancreatic cancer metastases

**DOI:** 10.1101/2022.08.27.504988

**Authors:** Ryan R. Kawalerski, Mariana Torrente Gonçalves, Chun-Hao Pan, Robert Tseng, Lucia Roa-Peña, Cindy V. Leiton, Luke A. Torre-Healy, Taryn Boyle, Sumedha Chowdhury, Natasha T. Snider, Kenneth R. Shroyer, Luisa F. Escobar-Hoyos

## Abstract

Keratin 17 (K17), an oncofetal intermediate filament protein, is one of the most abundantly expressed proteins in pancreatic ductal adenocarcinomas (PDACs) of the most aggressive molecular subtype. The mechanistic roles of this protein in malignancy, however, are largely unexplored. Here we show that K17 expression and disassembly enhances tumor growth and metastatic potential and shortens survival. Using mass spectrometry in K17 isolated from patient’s tumors, we identified a hotspot phosphorylation site in serines 10-13. Site-mutagenesis revealed that phosphorylation of this hotspot is sufficient to disassemble K17 and promote its nuclear translocation. *In silico* and pharmacologic inhibition studies uncovered the role of the PKC/MEK/RSK pathway in the phosphorylation and disassembly of K17. Murine models bearing tumors expressing phosphomimetic mutations at the serine hotspot displayed enhanced metastases, compared to mice bearing tumors expressing wild-type K17 or phosphorylation-resistant K17. Lastly, we found that detergent-soluble nuclear K17 promotes the expression of metastasis promoting genes in both patient and murine tumors. These results suggest that phosphorylation at specific serines is sufficient to promote pancreatic cancer metastasis and shorter survival, and that these sites could provide novel, druggable therapeutic domains to enhance PDAC patient survival.

## Introduction

Pancreatic ductal adenocarcinoma (PDAC) is a highly aggressive cancer, with a 5-year survival rate of only 10% due to propensity for metastatic dissemination and intrinsic resistance to first-line chemotherapies (Kleeff et al. 2016; Chu et al. 2017; Yao et al. 2020). The clinical management of PDAC is still limited by lack of therapeutic options, tumor heterogeneity that limit candidacy for targeted therapies, and the void in our knowledge of tumor-specific features to guide the development of novel therapeutics (Moffitt et al. 2015; Bailey et al. 2016; Aguirre et al. 2018; Aung et al. 2018; Pishvaian and Petricoin 2018; Maurer et al. 2019). There is therefore an urgent need to identify therapeutic targets and develop biomarker-based targeted approaches for PDAC treatment.

K17 is a type I embryonic keratin expressed broadly in ectodermal stem cells during embryogenesis, and its expression is suppressed in most adult somatic tissues (McGowan and Coulombe 1998b). In carcinomas of the lung, pancreas, cervix, and other sites, K17 is re-expressed for reasons that are not fully understood (Real et al. 1993; Mikami et al. 2015; Roa-Pena et al. 2019; Roa-Pena et al. 2021). In PDAC, K17 expression is a hallmark of the basal-like subtype, which is associated with the shortest patient survival and resistance to first-line therapies (Wang et al. 2011; Grasso et al. 2017; Aung et al. 2018; Roa-Pena et al. 2019; Pan et al. 2020; Roa-Pena et al. 2021). Growing evidence, however, suggests that K17 expression contributes to a variety of cancer hallmarks, suggesting its role as a potent oncoprotein (Ide et al. 2012; Mikami et al. 2015; Wu et al. 2017; Liu et al. 2018; Wang et al. 2019; Liu et al. 2020; Baraks et al. 2021).

As part of cellular homeostasis, keratins form stable supercoils that integrate into a larger structural network. Intermediate filament networks are repeatedly dissociated, or “disassembled,” from larger filamentous structures in a perinuclear distribution, after which they traffic outwardly for reincorporation into the network at the cell’s periphery (Windoffer et al. 2011; Moch et al. 2013). Although keratins were once thought to localize exclusively in the cytoplasm (Hobbs et al. 2016), we and others reported that K17 demonstrates nuclear localization to promote tumor growth by driving cell proliferation and altering gene expression (Escobar-Hoyos et al. 2015; Hobbs et al. 2015). The processes through which K17 assumes this non-canonical localization, though likely after disassembly from its filament network, remains largely unknown.

Post-translational modifications (PTMs) are regulators of the functions of intermediate filaments (Snider and Omary 2014). Among these, phosphorylation, sumoylation, and acetylation promote keratin disassembly, with phosphorylation accounting for the majority of PTMs found on keratins across normal and malignant tissues (Snider and Omary 2014; Hornbeck et al. 2015). Importantly, hyperphosphorylation of keratins directly increases cancer cell migration and metastatic capacity *in vitro* and *in vivo*, respectively, by re-arranging the cytosketon (Celis et al. 1983; Busch et al. 2012; Holle et al. 2017). Thus, we hypothesized that identification of a disassembly-driving phosphorylation site on K17, as well as site-specific phosphorylation inhibition, may suppress its oncogenic capacity by disrupting K17 nuclear localization and suppressing its pro-tumorigenic functions. Identification of PTMs that drive K17-disassembly may thus present an opportunity for therapeutic intervention in K17-expressing cancers.

We report novel phosphorylation sites in K17 that are sufficient to disassemble K17 from its filamentous network. Chemical approaches identified the role of PKC/MEK/RSK pathway in promoting these phospho-sites and subsequent K17-disassembly. *In vivo* studies in pre-clinical mouse models of PDAC found that phosphorylated and nuclear detergent-soluble K17 promotes PDAC cell migration and metastasis by enhancing the expression of metastasis-promoting genes. These results suggest that targeting K17’s disassembly represents a therapeutic opportunity to decrease metastatic dissemination and extend patient survival.

## Results

### K17 expression and disassembly enhances pancreatic cancer aggressive features

To assess the impact of K17 on PDAC tumor dynamics, we generated a K17 gain-of function model using a murine PDAC cell line (derived from the KPC mouse model), which does not endogenously express K17 protein. Isogenic cells, either expressing K17 or EV control, were assessed for K17 expression (**Fig. 1A-B**). We orthotopically implanted EV or K17 KPC cells into the pancreata of syngeneic mice. Mice with K17-expressing tumors had shorter survival compared to mice bearing tumors lacking K17 (HR = 4.373, *P*<0.0001, **Fig. 1C**). Furthermore, K17-expressing tumors were twice as large and retained K17 expression by the time of death (**Fig. 1D**). Importantly, K17 tumors exhibited enhanced hallmarks of tumor aggression, as demonstrated by larger relative areas of tumor necrosis (**Fig. 1E**) and increased incidence in hepatic metastasis compared to EV tumors (**Fig. 1F**). Together, these results suggest a cell-intrinsic mechanism by which K17 triggers three key hallmarks of PDAC aggression: growth, necrosis, and metastasis.

**Figure 1.**
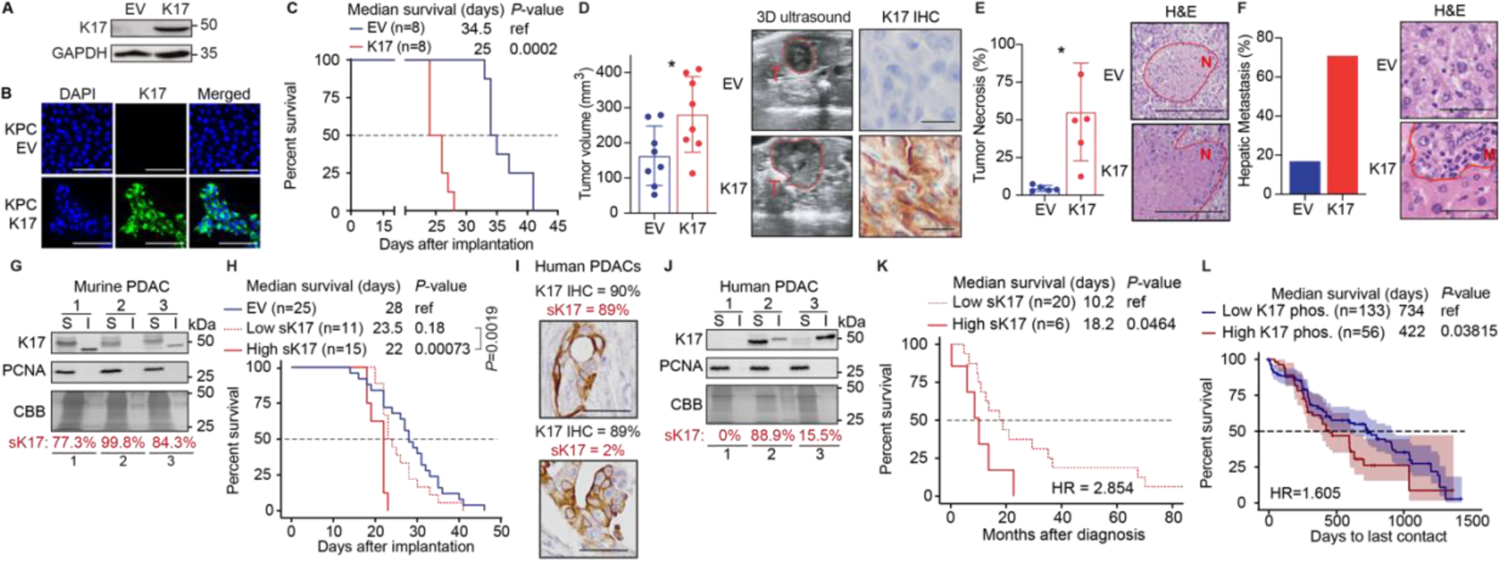
Keratin 17 enhances tumor aggression in a murine orthotopic model of PDAC and detergent-soluble Keratin 17 is negatively prognostic in human and murine PDACs. **A.** Immunoblot of lentiviral-generated KPC cells transduced to express K17 or empty vector (EV) for orthotopic implantation into the pancreata of immunocompetent C57BL/6 mice. **B.** Immunofluorescent labeling of K17 protein in lentiviral-generated clonal KPC cells transduced to express K17 or EV. K17 is shown in green, and the nucleus in blue (DAPI). Scale = 100μm. **C.** Kaplan-Meier curves show the overall survival in mice implanted with EV or K17-expressing PDACs. Significance determined using Log-rank Mantel–Cox test. **D.** Tumor size measured by 3D ultrasound. K17 tumors were on average larger than EV tumors, determined by ultrasound imaging. Representative IHC images indicating that K17 tumor cells implanted into mice retain expression of K17 *in vivo*. Scale = 20μm. Data are mean ± *s.d, n* = 8 mice per group. Student’s *t*-test. **E.** Percent of tumor necrosis in tumors. K17 overexpression increases the extent of tumor necrosis *in vivo*, representative H&E images. Scale = 200μm. Data are mean ± *s.d, n* = 5 mice per group. Student’s *t*-test. **F.** Percentage of hepatic metastases. K17 overexpression increases metastasis, representative H&E images. Scale = 50μm. Data are percentages*, n* = 5 mice per group. **G.** Western blot of detergent soluble (S) and detergent-insoluble (I) fractions of K17 from isogenic tumors with K17 overexpression. Molecular weight shift of K17 may be due to the hyperphosphorylation in the detergent-soluble fraction. *n* = 3. **H.** Kaplan-Meier curves demonstrate overall survival of mice with absent K17 (EV), low detergent-soluble-K17 and high detergent-soluble-K17 PDACs. Significance determined using Log-rank Mantel–Cox test. **I.** Representative images of K17 IHC of human PDACs, demonstrating that K17 disassembly is independent of K17 IHC score. Scale = 50μm. **J.** Representative blots of detergent soluble (S) and detergent-insoluble (I) fractions from human PDACs, showing a range of K17 disassembly. *n* = 3. **K.** Kaplan-Meier curves show the overall survival of patients with low detergent-soluble-K17 and high detergent-soluble-K17 PDACs. Significance determined using Log-rank Mantel–Cox test. **L.** Kaplan-Meier curves of PDAC cases from the Clinical Proteomic Tumor Analysis Consortium (CPTAC) stratified by either low or high K17-phosphorylation status. Significance determined using Log-rank Mantel–Cox test. *s.d,* standard deviation; *n*, number of repetitions. Coomassie Brilliant Blue (CBB). Hazard Ratio (HR). \**P*< 0.05; \*\**P*< 0.01; \*\*\**P*< 0.001.

As keratin subcellular reorganization requires disassembly and nuclear K17 regulates tumor cell functions (Escobar-Hoyos et al. 2015; Hobbs et al. 2015), we sought to investigate the association between K17-disassembly and PDAC aggression by analyzing patient and murine survival. We first isolated detergent-soluble and detergent-insoluble K17 fraction using high-salt extraction from a subset of K17 KPC murine PDACs to assess K17 disassembly by western blot. Analyses revealed a range of detergent-soluble K17 (sK17) expression across these isogenic tumors (77.3%-99.8%) (**Fig. 1G**). The fraction of sK17 in these tumors strongly correlated with mouse survival, with tumors in the highest quartile of sK17 (high sK17) associated with shorter survival than those outside of the highest quartile (low sK17) (**Fig. 1H**). This suggests that K17-disassembly is a prerequisite to engage the oncogenic roles of K17 in PDAC. We next hypothesized that PDAC patients with tumors expressing similar levels of K17 protein could have differences in survival explained by the K17 disassembly status. We processed flash frozen human PDAC samples for detergent-soluble and detergent-insoluble extracts and found that PDACs expressing similar levels of total K17 staining by IHC and viable tumor area had variable levels of sK17 (**Fig. 1I-J**). Tumors with sK17 levels in the highest quartile were associated with decreased patient survival compared to other quartiles (**Fig. 1K**). While rapid cell-cycle progression contributed to keratin filament disassembly, *PCNA* expression did not correlate with sK17 in human and murine tumors (**Fig. 1J, Supplementary Fig. S1A-C**). Furthermore, analysis of high K17-expressing PDAC cases from the Clinical Proteomic Tumor Analysis Consortium (CPTAC) stratified by low or high K17-phosphorylation status, a proxy endpoint of keratin disassembly, demonstrated that those expressing high K17-phosphorylation correlated with a worse survival outcome versus low K17-phosphorylation cases (**Fig 1L**). Overall, these data suggest that the mechanisms that regulate K17-disassembly may impact the survival of PDAC patients.

### Serine phosphorylation (S10-13) drives K17 disassembly

The functional properties of keratins are largely regulated by post-translational modifications (PTMs), of which phosphorylation is critical for reorganization of keratin filament networks via cytoplasmic and perinuclear disassembly (Snider and Omary 2014). To identify the PTMs on K17 that may enable its disassembly, we performed liquid chromatography-mass spectrometry (LC-MS) in K17 isolated from archival flash-frozen patient derived non-treated PDACs with high K17 expression (**Fig. 2A, Supplementary Fig. S2A**). LC-MS results showed a coverage of 97% of K17 residues and revealed phosphorylation of serines within the span of the 50 most N-terminal amino acids in all three patient samples (**Fig. 2B**), including previously unreported phosphorylation at serines 11, 12, 13. Protein sequence analysis of the K17 polypeptide (Blom et al. 1999; Blom et al. 2004) revealed that the K17 N-terminal S10-13 region is well-conserved across several species and across human keratin proteins in the type I keratin family (**Fig. 2C, Supplementary Fig. S2A**). In addition, we confirmed the previously reported K17-phosphorylation site at serine 44 (Ser44) and using a previously reported antibody (Pan et al. 2011), found that K17-phosphorylation at S44 is highly enriched in detergent-soluble fractions of K17 (**Supplementary Fig. S2B**). Overall, these findings suggest that phosphorylation, particularly on N-terminal serine residues, may regulate K17-disassembly in human PDACs.

**Figure 2.**
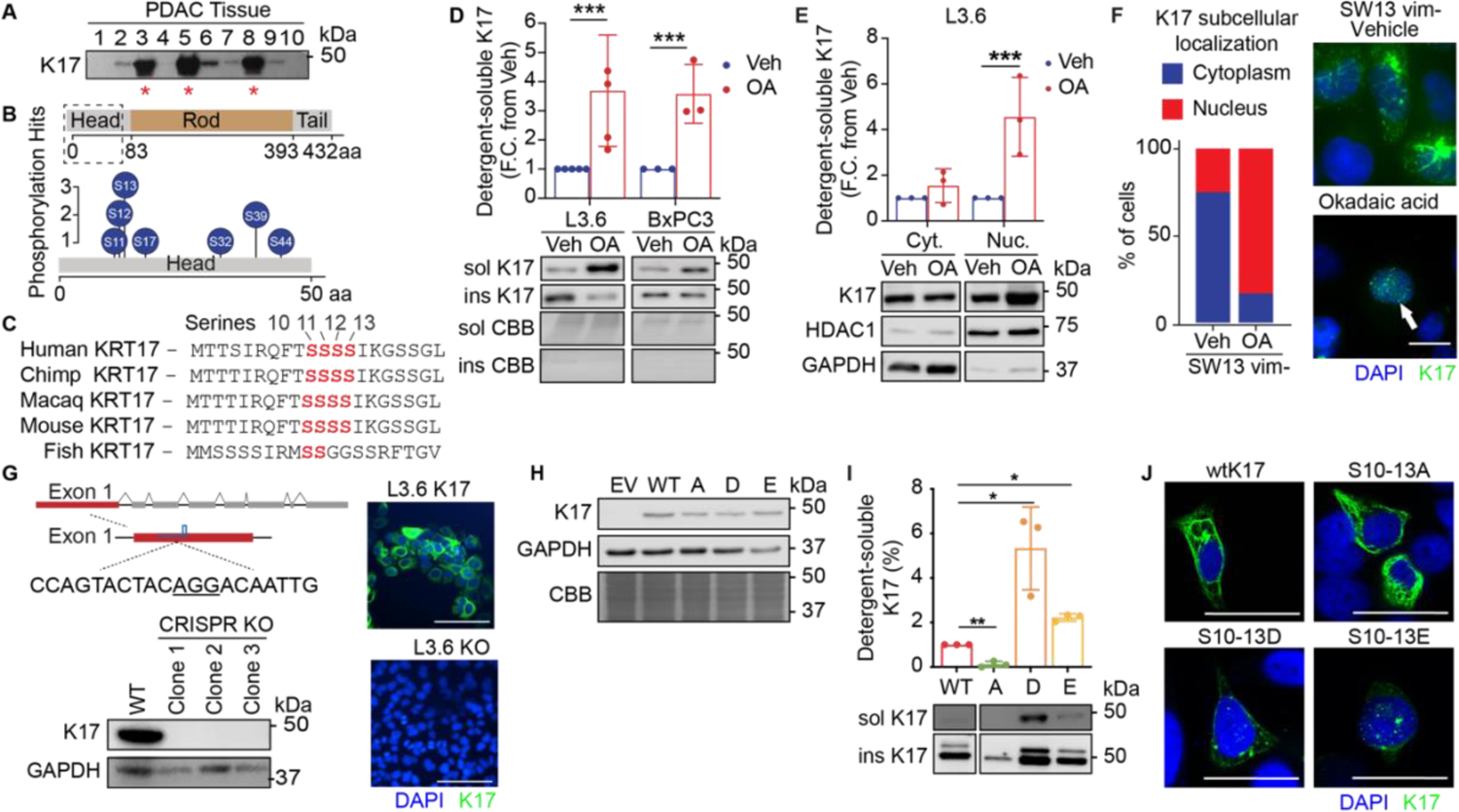
K17 is differentially phosphorylated in its detergent-soluble state, and N-terminal serine phosphorylation promotes disassembly. **A.** Western blot of human PDAC samples showing three high-K17 expressing tumors chosen for mass spectrometry (indicated with a red *). Equal micrograms of protein were loaded for each sample. **B.** Illustration of results from liquid chromatography mass spectroscopy of the three PDACs from (A), which identified serine phosphorylation on the N-terminal residues indicated (head of the keratin). **C.** Alignment of K17 polypeptide in different vertebrate species show identification of a conserved serine hotspot (Ser10-13). **D.** Western blot depicting detergent-soluble and detergent-insoluble K17 after treatment with okadaic acid (OA). Treatment leads to disassembly of K17 in L3.6 and BxPC3 PDAC cell lines, compared to vehicle control (Veh), shown by fold change (FC). Data are mean ± *s.d, n* = 3 independent repetitions. **E.** Western blot depicting detergent-soluble and detergent-insoluble K17 in cytoplasm (Cyt.) and nuclei (Nuc.) after treatment with okadaic acid (OA). K17 is preferably localized to the nucleus upon treatment of PDAC cells with okadaic acid, as shown by fold change of total K17 level. Data are mean ± *s.d, n* = 3 independent repetitions. **F.** Subcellular localization of K17 after okadaic acid (OA) treatment by immunofluorescence in SW13 vim-cells. Scale =5μm. 200 cells were quantified in each condition. **G.** Illustration of K17 edited by CRISPR-Cas9 technique used to generate K17 knockout (KO) in L3.6 cells. Western blot validation of the knockout in three L3.6 cell clones and immunofluorescence images of L3.6 expressing K17 and the KO clone. Filament form of K17 is shown in green, and the nucleus was stained by DAPI, in blue. Scale = 100 μm. **H.** Western blot depicting total K17 expression following lentiviral transfection of K17 phospho-mutants into L3.6 clone cells. **I.** Percent Detergent-soluble K17 in functional phospho-K17 mutants in original knockout cells. S10-13 hotspot is sufficient to regulate K17 disassembly, as shown by fold change in detergent-soluble K17 level. Data are mean ± *s.d, n* = 3 independent repetitions. **J.** Immunofluorescence of functional phospho-K17 mutants shows that L3.6 gain-of-function S10-13D and S1013-E K17 mutants have a distinctly punctate, not filamentous, cytoplasmic K17 organizational pattern. Filament and punctate forms of K17 are shown in green, and the nucleus was stained by DAPI, in blue. Scale = 20 μm. *s.d,* standard deviation; *n*, number of repetitions. Coomassie Brilliant Blue (CBB). \**P*< 0.05; \*\**P*< 0.01; \*\*\**P*< 0.001.

To determine whether phosphorylation at the identified N-terminal serine residues is required for K17-disassembly, we induce keratin-network disassembly using broad-spectrum modulators of phosphorylation in human PDAC cell lines that endogenously express K17. Inhibition of the phosphoserine/threonine phosphatase 1 and 2a (PTP-1/2a) by okadaic acid (OA) at concentrations previously identified to modulate inhibit de-phosphorylation in intermediate filaments (Snider and Omary 2016),inhibited K17 disassembly in PDAC cells more than 3-fold compared to cells treated with vehicle control (**Fig. 2D).** This enhanced disassembly correlated with K17 localization to the nucleus in human PDAC cells (**Fig. 2E, Supplementary Fig. S2C-E**). To validate these findings, and to test if K17 disassembly is dependent on the disassembly of other intermediate filaments, we engineered the SW13 VIM negative cell line, a small cell adrenal cortical carcinoma line that lacks keratins and other intermediate filament proteins, to transiently express K17. We found that upon OA treatment, detergent-soluble K17 was increased and was found mainly in the nuclei of cells, consistent with previously observed K17-mediated disassembly effects (**Fig. 2F**). This supports prior findings that non-filamentous K17 acts as a nuclear protein in the context of cancer (Escobar-Hoyos et al. 2015). To test whether tyrosine phosphorylation contributes to disassembly, we used a competitive inhibitor of phosphotyrosyl protein phosphatases (sodium orthovanadate, OV), at a concentration shown to modulate IFs(Strnad et al. 2002). Treatment with OV showed limited changes in disassembly (**Supplementary Fig. S2F**). Thus, phosphorylation at serines, rather than tyrosines likely provides a more biologically potent and actionable mechanism to disassemble K17 in PDAC.

Serines 10-13 are located in the N-terminal region of K17, a domain that is frequently phosphorylated on other keratins (Omary et al. 2006). Having demonstrated that these residues are phosphorylated on K17 isolated from PDAC specimens, we next sought to determine if phosphorylation of this region was independently sufficient for K17-disassembly. Using the L3.6 human PDAC cell line that endogenously expresses K17 (Gysin et al. 2005b), we generated three CRISPR-mediated K17 knockout clones (**Fig. 2G**). In each of the three clones, we re-expressed either wild-type K17 (wtK17) or K17 with mutations in S10-13 in three forms to modulate phosphorylation: 1. alanine, loss-of-function (S10-13A), 2. aspartate, gain-of-function (S10-13D), and 3. glutamate, gain-of-function (S10-13E) (**Fig. 2H**). Both aspartate and glutamate are phosphomimics for serine phosphorylation. The generated cell lines from each clone were used as biological replicates for *in vitro* experiments to evaluate how phosphorylation at the serine-rich site impacts K17 disassembly. Analysis of K17 disassembly for each generated cell line showed that the loss-of-function S10-13A mutant resulted in an approximately 10-fold decrease in disassembly compared to wtK17. Importantly, both gain-of-function mutants, S10-13D and S10-13E, exhibited increased disassembly compared to wtK17 (**Fig. 2I**). Confocal imaging of these cells showed that both the wtK17 and S10-13A models had K17-filamentous networks, while the S10-13D and S10-13E models displayed a punctate pattern with increased K17 localization in the cell nuclei, as seen with OA treatment (**Fig. 2J**). These results suggest that K17-phosphorylation at serine 10-13 is necessary and sufficient for its disassembly.

### A PKC/MEK/RSK signaling cascade mediates K17 disassembly

To identify the kinase(s) responsible for K17-phosphorylation at the S10-13 hotspot, we first performed bioinformatic predictions using NetPhos (Blom et al. 2004), GPS 5.0 (Wang et al. 2020) and other methods (Safaei et al. 2011) and identified that S10-13 is likely phosphorylated by PKC-driven mechanisms (**Fig. 3A**). Stimulation with a PKC activator, 12-O-Tetradecanoylphorbol-13-acetate (TPA, also known as PMA, a potent phorbol ester mimicking diacylglycerol), induced time-dependent K17-disassembly (**Fig. 3B**) of wtK17 but not the S10-13A mutant, indicating that the TPA-mediated signal is specific to S10-13 (**Fig. 3C**). We next evaluated whether PKC kinases themselves or downstream kinases MEK1/2 or RSK were responsible for mediating phosphorylation. RSK, which comprises 4 isoforms (RSK1-4) (Casalvieri et al. 2017), has been reported to phosphorylate K17 on S44, although silencing RSK1 in murine keratinocytes does not impact TPA-induced phosphorylation of S44 on K17 (Pan et al. 2011). We thus performed co-treatment experiments whereby cells were serum-starved for 24h, incubated with inhibitors of PKC or known downstream kinases for 1h, and then treated with TPA for 1h to activate the PKC pathway. Treatment with small molecule inhibitors of either PKC alpha or beta isoforms (Go6976), MEK1/2 kinases (Selumetinib), or p90 RSK family (BI-D1870) kinases blocked TPA induced K17-disassembly (**Fig. 3D-F**). All treatments were conducted at concentrations below IC50. The largest differences in sK17 were seen after treatment with BI-D1870, suggesting that the p90 RSK proteins, which constitute the last kinase activation step of the PKC signaling cascade prior to IF phosphorylation, are the most likely candidate kinases that phosphorylate K17. Thus, a PKC/MEK/RSK pathway phosphorylates K17 and targeting this event may mitigate K17-driven properties of tumor aggression (**Fig. 3G**).

**Figure 3.**
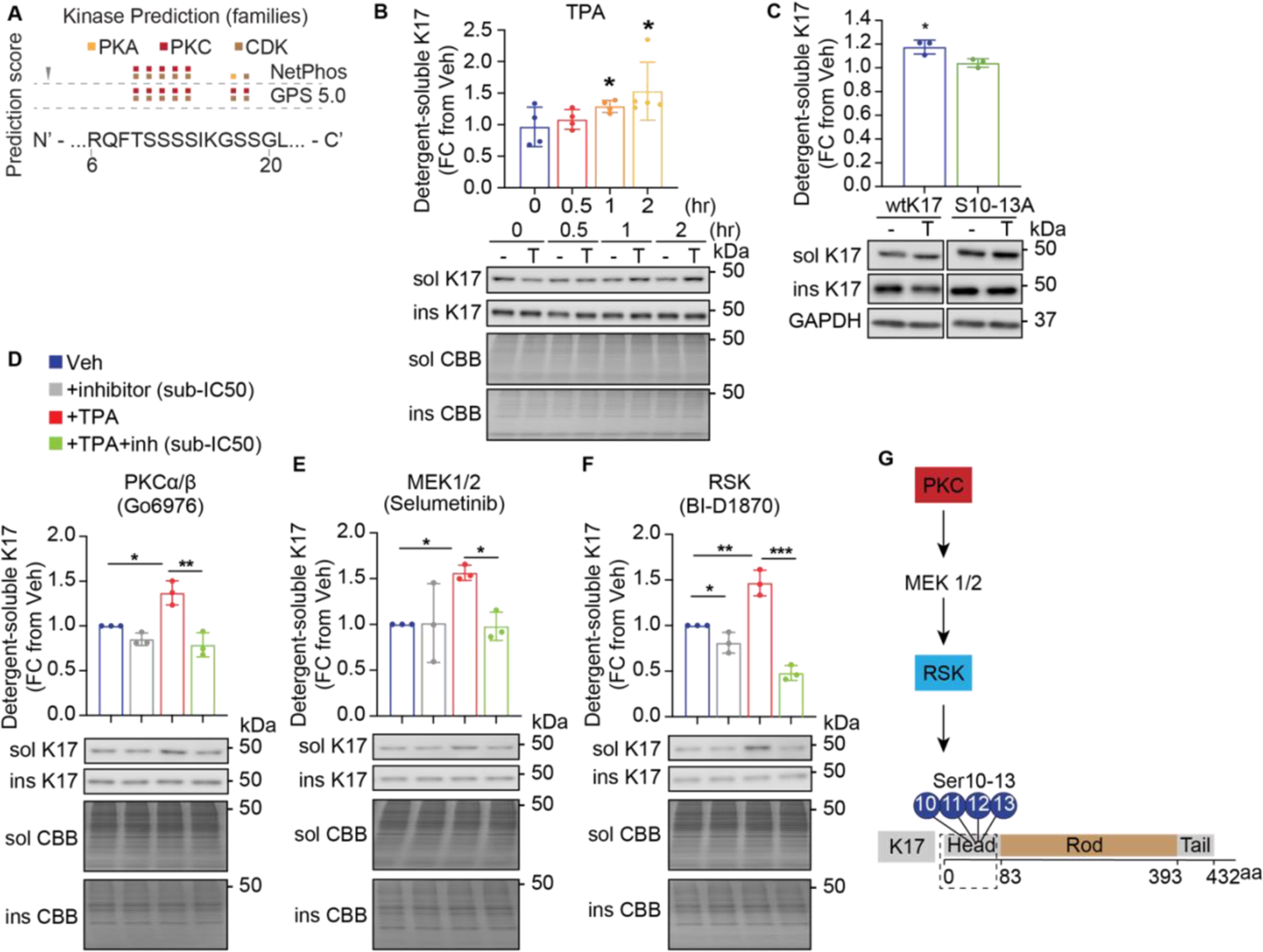
K17 is phosphorylated by a PKC-mediated signaling cascade. **A.** Bioinformatic prediction from NetPhods and GPS 5.0, for serine, threonine, or tyrosine phosphorylation in K17 indicates that the S10-13 region is likely to be phosphorylated by a PKC-mediated pathway. **B.** Western blot of detergent-soluble and detergent-insoluble K17 in L3.6 cells with TPA treatment (T, 400nM) strongly induces K17-disassembly after 1h, shown by fold change of detergent-soluble K17. Data are mean ± *s.d, n* = 5 independent repetitions. Student’s *t*-test. **C.** Western blot of detergent-soluble and detergent-insoluble K17 in L3.6 cells with TPA treatment (T, 400nM) in loss of function mutant and wt K17. Data are mean ± *s.d, n* = 3 independent repetitions. Student’s *t*-test. Data are mean ± *s.d, n* = 3 independent repetitions. Student’s *t*-test. **D-F.** Treatment of L3.6 cells with sub-IC50 doses of inhibitors for PKC (**D**), MEK1/2 (**E**), and RSK (**F**) kinases for 1h, followed by 1h-TPA stimulation, reduced the levels of detergent-soluble K17, shown by fold change of detergent-soluble K17 and western blot. Data are mean ± *s.d, n* = 3 independent repetitions. Student’s *t*-test. **G.** Illustration of the PKC and downstream kinases proposed to phosphorylate K17 at the N-terminal S10-13 region. *s.d,* standard deviation; *n*, number of repetitions. Coomassie Brilliant Blue (CBB), detergent-insoluble (insol), detergent-soluble (sol), 12-O-Tetradecanoylphorbol-13-acetate (TPA). \**P*<0.05, \*\**P*<0.01, \*\*\**P*<0.001.

### Detergent-soluble K17 enhances pancreatic cancer metastasis by regulating gene expression programs

To assess the role of the K17 S10-13 phosphorylation on tumor dynamics, we orthotopically implanted L3.6 cells stably expressing either wt-K17 or phospho-mutated K17 into the pancreata of immunodeficient NU/J mice. Mice bearing tumors expressing loss-of-phosphorylation (LOP) K17 (S10-13A) survived longer than those expressing gain-of-phosphorylation (GOP) K17 mutants (S10-13D or S10-13E) (**Fig. 4A**). Analysis of sK17 levels in tumors confirmed differences in K17 disassembly and nuclear localization between the LOP versus the GOP mutant (**Fig. 4B, Supplementary Fig. S3A**). In addition to differences in survival, GOP K17 tumors were larger, more necrotic and had more hepatic metastases compared to S10-13A K17 tumors (**Fig. 4C-D, Supplementary Fig. S3C**). In agreement, *in vitro* cell proliferation and wound closure assays demonstrated that LOP had decreased cell numbers and migration, compared to WT and GOP mutants (**Supplementary Fig. 3C-E**).

**Figure 4.**
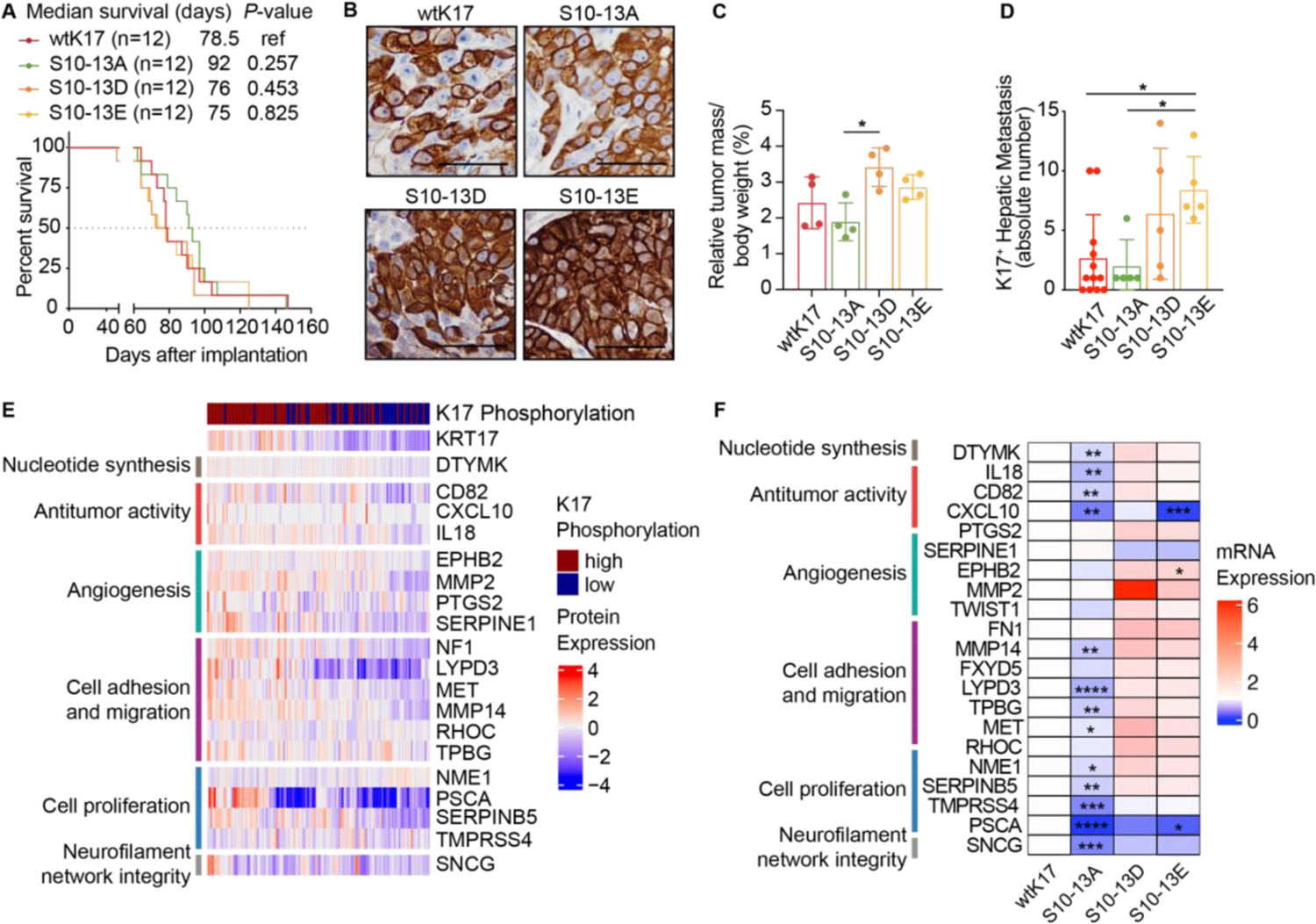
K17-phosphorylation impacts tumor growth and metastases *in vivo*. **A.** Kaplan-Meier curves for immunodeficient NU/J mice implanted orthotopically with L3.6 cells stably expressing K17 with loss of phosphorylation function (LOPF), S10-13A. Significance determined using Log-rank Mantel–Cox test. **B.** Representative images of K17 IHC in murine tumors expressing wtK17 or phospho-mutant K17. Scale = 100μm. **C.** Relative tumor mass. Tumors of immunodeficient NU/J mice implanted orthotopically with L3.6 cells stably expressing gain of phosphorylation function (GOPF) S10-13D K17 are larger than those expressing loss of phosphorylation function S10-13A, as determined by tumor mass relative to body weight. Data are mean ± *s.d, n* = 4 mice per group. Student’s *t*-test. **D.** K17-positive hepatic metastasis assessed by IHC. Mice implanted with L3.6 cells expressing S10-13D or S10-13E K17 have more hepatic metastasis. Data are mean ± *s.d, n* = 5-8 mice per group. Student’s *t*-test. **E.** Protein expression of metastasis-associated genes from CPTAC PDAC samples annotated with K17 protein expression and total K17-phosphorylation status. Expression expressed in fold change. **F.** qRT-PCR mRNA expression of metastasis-associated genes in L3.6 cells expressing phospho-K17 mutants, relative to wild-type (wt) K17. Data expressed as fold change of wt K17. Data are mean ± *s.d, n* = 3 repetitions. Student’s *t*-test. \**P*<0.05, \*\**P*<0.01, \*\*\**P*<0.001.

Given that sK17 mainly localizes to the nucleus of tumor cells (**Fig. 2**) and previous reports suggest that K17 binds to chromatin and regulates gene expression (Jacob et al. 2020), we hypothesized that sK17 promotes the expression of genes that impact metastatic capacity. Analysis of RNA and protein expression profiles of PDAC cases from the CPTAC patient cohort revealed that the expression of metastasis-promoting genes correlated with increased total K17-phosphorylation (**Fig. 4E**). To determine if K17 causes differential expression of metastatic genes, we performed RNA-seq in K17 proficient and deficient cells. By differential gene expression and gene enrichment analyses, we found that K17 expression strongly upregulated genes involved in metastasis (**Fig. 4F**). From these enriched genes, we selected the top 22 and analyzed their expression in L3.6 cells that expressed either wtK17 or S10-13 phospho-mutants. In the LOP S10-13A mutant, mRNA levels of *DTYMK*, *IL-18*, *CD82*, *CXCL10*, *MMP14*, *LYPD3*, *TPBG*, *MET*, *NME1*, *SERPINB5*, *TMPRSS4*, *PSCA* and *SNCG* were downregulated, relative to wt-K17 cells (**Fig. 4F**), while *EPHB2* was upregulated in S10-13D and S10-13E, relative to wt-K17 control. Altogether, these alterations suggest that K17 S10-13 phosphorylation affects the expression of genes that regulate anti-tumor activity related to inflammation, angiogenesis, cell adhesion, migration, and proliferation (**Fig. 4F**), resulting in enhanced tumor growth and metastatic spread.

## Discussion

Here, we show that phosphorylation drives the disassembly and nuclear localization of K17 in cancer, impacting metastasis and survival. While phosphorylation has been reported for other keratins and *in silico* analysis showed that phosphorylation levels of S13, S24, S32, and S39 were enhanced in several cancers compared to primary tissues (Li et al. 2021), our work is the first to demonstrate that phosphorylation of a conserved N-terminal, serine-rich domain in K17 increases the expression of metastasis-promoting genes. This mechanism contrasts to other reports that have linked keratins to metastatic capacity by cytoskeletal rearrangements in epithelial-to-mesenchymal transition (Seltmann et al. 2013). The identification of phosphorylation of a conserved serine-rich region of K17 derived from patient tumors, suggests that these PTMs are clinically relevant. Furthermore, we show that this serine-rich region is likely phosphorylated by a PKC/MEK/RSK pathway, which suggests a therapeutic opportunity. In summary, this is the first report that K17-phosphorylation is linked to pancreatic cancer aggression.

Our *in vivo* data revealed that K17 expression is an independent determinant of pancreatic cancer aggression, as seen in animal models bearing K17-expressing PDAC. These findings provide a biological underpinning to prior patient gene expression analyses (Moffitt et al. 2015; Bailey et al. 2016; Maurer et al. 2019) that correlate with patient prognosis and disease aggression (Hayashi et al. 2020). Importantly, K17 expression is a hallmark and biomarker to identify the most aggressive molecular subtype of PDAC (Collisson et al. 2011; Moffitt et al. 2015; Bailey et al. 2016; Maurer et al. 2019; Hayashi et al. 2020), however, the mechanisms through which K17 drives PDAC aggression are not fully understood. We and others previously reported that K17 promotes tumor growth upon nuclear localization in cervical squamous cell carcinoma and premalignant epidermal keratinocytes (Escobar-Hoyos et al. 2015; Hobbs et al. 2015). Disassembly of K17 likely precedes nuclear localization, and therefore may be required for oncogenesis. However, little is yet known about the mechanisms of action that regulate K17 disassembly. Phosphorylation is a known driver of intermediate filament disassembly, including of type I and II keratins, and is modulated by several kinases that target Ser/Thr residues in the head, as identified in this report, or tail domains of the proteins (Omary et al. 2006; Snider and Omary 2014). Yet, no association has been established between phosphorylation and oncogenic functions of K17. Proteomic analysis previously identified phosphorylation sites in the C-terminus of K17, specifically in Y398 and Y410, and *in silico* analysis also suggested that phosphorylation at these sites regulate protein nuclear localization (Hobbs et al. 2016). Conversely, we discovered that phosphorylation on S10-13 of the N-terminal region is sufficient to impact disassembly and nuclear localization of K17 in PDAC.

Discovery of the S10-13 region as a phosphorylation site on K17 is consistent with other findings that N-terminal serines of keratins proteins are phosphorylated to drive intermediate filament disassembly in cancer (Ku et al. 1998; Ku et al. 2002; Ridge et al. 2005), including keratins 8 (S73) (Ku et al. 2002) and 18 (S33) (Ku et al. 1998; Karantza 2011). Disassembly of keratins, particularly keratin 8 (K8), a type II keratin, has also been demonstrated in pancreatic and gastric tumor cells and transformed amniotic cells to promote metastatic potential (Celis et al. 1983; Busch et al. 2012; Holle et al. 2017). Keratin phosphorylation may also disrupt phosphorylation of pro-apoptotic proteins, thus preventing activation of proteins by kinases active under stress (Ku and Omary 2006; Omary et al. 2006). In K8, phosphorylation of S73 is required for cellular apoptosis and S431 for enhanced migration of pancreatic cancer cells (Busch et al. 2012). However, in the context of type I keratins, including K17, there is limited evidence regarding disassembly-mediated metastasis. In K18, another type I keratin, phosphorylation of S52 is involved in filament reorganization (Busch et al. 2012), but in K17, phosphorylation of N-terminal S44 was required for cell growth in epithelial cells (Pan et al. 2011). While all previous reports link the phosphorylation and disassembly of keratins to metastasis through cytoskeletal reorganization, here we report for the first time that K17 phosphorylation and subsequent disassembly, lead to nuclear translocation and increased expression of metastasis-associated genes. This suggests that K17 disassembly leads to tumor aggression by changing the transcriptome of cells and by changing the physical properties of the intermediate filaments.

Prior work has shown that pancreatic cancers expressing high levels of K17 are resistant to commonly deployed first-line chemotherapeutic agents, notably gemcitabine (Wang et al. 2011; Aung et al. 2018; Pan et al. 2020; Yao et al. 2020). The finding that K17-phosphorylation at the N-terminus S10-13 is regulated by a PKC/MEK/RSK pathway is important because it suggests that K17 may be sensitive to targeted kinase inhibition. Accordingly, others have suggested that a pathway involving RSK likely phosphorylates K17 (Pan et al. 2011), and that PKC/MEK/RSK pathway may drive pancreatic tumor aggression and metastasis, both *in vitro* and *in vivo* (Boucher et al. 2000; Gysin et al. 2005a; Ma et al. 2011; Alagesan et al. 2015). Despite these findings, phase II clinical trials of pancreatic cancer patients treated with Selumetinib (MEK 1/2 inhibitor) failed to improve patient overall survival (Bodoky et al. 2012; Chung et al. 2017). Therefore, using K17 as a predictive biomarker for therapeutic response in pancreatic cancer could guide effective therapeutic intervention with other inhibitors, such as RSK inhibitors. In summary, our findings have identified a novel oncogenic mechanism for K17 in pancreatic cancer that has therapeutic implications; further study is required to explore these mechanisms as a therapeutic vulnerability of these deadly tumors.

## Materials and Methods

### Patient tissue acquisition and survival analysis

Analysis of detergent-soluble K17 was performed on 26 patient archived flash-frozen PDACs from a cohort originally described in Roa-Peña *et al*. The Studies were performed in accordance with guidelines and regulations of the Stony Brook Medicine Institutional Review Board protocol 94651. Stratification of samples by high salt protein extraction used for survival analyses was performed at the highest quartile as this cutoff corresponded to approximately 80% of the total K17 protein content isolated in the detergent-soluble fraction for both the murine and human samples.

### Murine orthotopic xenograft studies

All experimental procedures were approved by the Institutional Animal Care and Use Committee (IACUC) at Stony Brook University and are in accordance with the Guide for the Care and Use of Laboratory Animals from the National Institutes of Health.

#### Study I

Murine PDAC KPC cells with/without K17 expression were mixed in 1:1 matrigel:Opti-MEM® (ThermoFisher Scientific) solution to a final volume of 30μL holding 150,000 cells, and orthotopically implanted into tail of the pancreas of C57BL/6J mice. Tumor growth was measured by weekly 3D ultrasound (Vevo 3100 Preclinical Imaging System; FUJIFILM VisualSonics).

#### Study II

L3.6 cells stably expressing either wild-type K17 or phosphorylation mutants S10-13A, S10-13D, or S10-13E were prepared as described in Study I, implanted in NU/J mice and followed by weekly 3D ultrasound. Mice were euthanized at days 22 and 52 post-implantation and organs were harvested for histological assessment of metastasis. Pancreatic tumors were flash-frozen in liquid nitrogen upon collection.

### Immunohistochemistry and scoring method

Immunohistochemical staining and scoring of formalin-fixed, paraffin-embedded tissues were performed as previously described (Escobar-Hoyos et al. 2014). Following antigen retrieval, human and KPC murine tissues were incubated with mouse monoclonal KDx K17 antibody (KDx Diagnostics Inc., Campbell, CA) and NU/J murine tissues were incubated with rabbit polyclonal K17 antibody (McGowan and Coulombe 1998a; Depianto et al. 2010; Hobbs et al. 2015) (kindly provided by Dr. Pierre Coulombe). Following primary antibody incubation, biotinylated anti-mouse IgG secondary antibodies (R.T.U. Vectastain ABC kit; Vector Laboratories, Burlingame, CA) or horse anti-rabbit IgG secondary antibodies (Vector Laboratories, Burlingame, CA) were added. Staining was developed using 3,3′-diaminobenzidine (DAB; Dako, Carpinteria, CA) followed by counter-staining with hematoxylin. K17 scoring was classified by one pathologist (KRS) based on a subjective assessment of absent (0), light (+1), or strong (+2) staining. The overall proportion of tumor cells with +2 intensity staining (the PathSQ score) was determined by blinded review of representative immunostained histologic sections from each case.

### Immunofluorescent imaging

Cells on glass slides were fixed in 95:5 solution of ethanol:acetate, followed by washing thrice with ice-chilled 1x Dulbecco’s Phosphate Buffered Saline (DPBS). Cells were permeabilized with 0.25% Triton X-100 in DPBS and blocked in 10% donkey serum in DPBS. Slides were incubated with Primary KDx K17 antibody, washed with DPBS, incubated with Alexa Fluor® goat anti-rabbit secondary antibody (Life Technologies, Carlsbad, CA), washed with DPBS and subsequent incubation with 5µg/mL 4′,6-diamidino-2-phenylindole (DAPI). Cells were mounted with Vectashield (Vector laboratories, Burlingame, CA) and imaged using an upright Leica DM 6000 confocal microscope.

### Mass spectrometry

Ten randomly selected flash-frozen human PDAC tissues were processed using the total protein extraction method as described below. Three tumors with the highest K17 protein expression were selected for high salt extraction. Detergent-insoluble fractions were loaded and adjusted for equal protein concentration in SDS polyacrylamide gels and stained by Coomassie Blue to confirm enrichment of protein fractions. The band corresponding to K17 was excised from the gel, and gel strips were digested with trypsin. For mass spectroscopy, peptides obtained were resolved on a nanocapillary reverse phase column using a 1% acetic acid/acetonitrile gradient at 300 nL/minute. Peptides were introduced into a Orbitrap Fusion tribrid (ThermoFisher Scientific) mass spectrometer and analyzed at 60K resolution. Results were analyzed for the identification of PTMs using Scaffold v.4.11.1.

### Cell culture

Murine PDAC cells expressing mutant Kras and Trp53 (derived from the Kras^G12D^; Tp53^R172H^; Pdx1-Cre25: KPC model), were a gift from Dr. Gerardo McKenzie, University of California San Diego. The human L3.6 PDAC cell line (KRAS^G12A^) was provided by Dr. Wei-Xing Zong (Rutgers University). The BxPC3 cell line was obtained from American Type Culture Collection. Cells were cultured in Dulbecco’s Modified Eagle Medium (DMEM, Gibco) supplemented with 10% fetal bovine serum (FBS, ThermoFisher Scientific) and 1% penicillin and streptomycin (P/S, Gibco), at 37°C in a humidified incubator under 5% CO_2_.

### Pharmacologic treatment

K17-disassembly following hyperphosphorylation was assessed using okadaic acid (OA, Cayman Chemical) and sodium orthovanadate (OV, Millipore Sigma). L3.6 and BxPC3 were grown to approximately 70% confluence in 10% FBS DMEM before treatment with OA (1µM) or OV (10mM). After treatment (40 minutes for OA, and 60 minutes for OV), cells developed a rounded morphology and cell lysate was collected. To assess inhibition of K17-phosphorylation via small molecular inhibitors, L3.6 cells were grown to 60% confluence, followed by media change to DMEM without FBS for 48h. Subsequently, kinase inhibitors for PKCα/β (Go6976, 10µM), MEK1/2 (Selumetinib, 2 µM) or RSK (BI-D1870, 20 µM) were added to fresh DMEM without FBS 1h before treatment with 12-O-Tetradecanoylphorbol-13-acetate (TPA, 400nM, Cayman Chemical), after which cell lysates were collected.

### Stable K17 overexpression in murine KPC cell line

KPC cells were transduced with either an empty vector (EV) pLVX-IRES-mCherry (Clontech, Mountain View, CA) or the same backbone vector expressing the human K17 open reading frame cDNA. mCherry-expressing cells were selected by fluorescence-activated cell-sorting, where the top 10% of cells were expanded in culture. Cell populations were tested for K17 expression by immunofluorescence and western blotting.

### Generation of CRISPR Cas9-mediated K17 knockout cell line

A CRISPR Cas9-mediated K17 knockout cell pool was generated in L3.6 cells by Synthego Corporation (Redwood City, CA, USA). Ribonucleoproteins containing the Cas9 protein and single guide RNA (CCAGTACTACAGGACAATTG) were electroporated into cells (https://www.synthego.com/resources/all/protocols). The genetic editing efficiency was assessed upon recovery 48h post-electroporation. Genomic DNA was extracted from the cells, PCR amplified, and Sanger sequenced and analyzed (ice.synthego.com).

### Stable K17 overexpression of K17 in L3.6 cell line

L3.6 cells were transduced with either an EV pLVX-IRES-Puro (Clontech, Mountain View, CA) or the same backbone vector expressing human K17 cDNA (wild-type or mutations of S10-13A, S10-13D, or S10-13E). Briefly, virus was collected from the centrifuged supernatant (DMEM with 1% FBS and without antibiotic) of HEK293T cells transfected with the above plasmids and frozen at −80°C until used. Supernatant with virus was thawed to transduce three L3.6 K17-knockout pooled population lines. After transduction, media was replaced with 2ug/mL puromycin for a 7-day selection of infected cells before returning to regular media. Cell populations were tested for K17 expression by western blotting.

### Transient transfection of L3.6 K17 knockout cells

Transient transfection with the pLVX-IRES-Puro plasmids, as mentioned above, was performed using Lipofectamine 3000 reagent (ThermoFisher Scientific). Cells were grown to 70% confluence, and media was switched from 10% FBS DMEM to 1% FBS DMEM 1h before transfection. Transfection reagent was placed on cell media for 24h before cells were lysed to assess K17 protein expression and disassembly.

### Total protein extraction

Total protein was extracted using radioimmunoprecipitation (RIPA) Lysis and Extraction Buffer (ThermoFisher Scientific) mixed with Halt™ Protease Inhibitor Cocktail (ThermoFisher Scientific). Lysates were sonicated briefly in pulses and centrifuged before supernatant was collected.

### Nuclear and cytoplasmic fractionation

Cells were fractionated into cytoplasmic and nuclear fractions using the NE-PER™ Nuclear and Cytoplasmic Extraction Reagents (ThermoFisher Scientific) according to the product manual.

### Western blot analysis

Cell lysate protein concentrations were measured using the DC™ Protein Assay kit (Bio-Rad). Equal amounts of protein were separated by a 10% SDS polyacrylamide gel electrophoresis. Immunoblotting was performed with primary antibodies to K17 (KDx) and GAPDH (Cell Signaling Technology), followed by infrared IRDye® goat anti-mouse or anti-rabbit IgG secondary antibodies (LI-COR Inc.). Western blot images were captured using the LI-COR Odyssey® Fc Imaging System and images were quantified using Image Studio Lite™ software (LI-COR Inc.).

### High-salt protein extraction

High-salt extraction for IF proteins was described previously (Snider and Omary 2016). Briefly, human and murine tissues or monolayer adherent cells were suspended in ice-cold Triton-X buffer (TXB) (1% Triton-X 100, 5mM EDTA in 1x PBS) containing protease and phosphatase inhibitors (PPI; ThermoFisher Scientific). Flash-frozen tissues were immediately placed in TXB after removal from storage and homogenized via Potter-Elvehjem PTFE pestle and glass tube. Cells were collected with ice-cold DPBS wash followed by scraping and resuspension into TXB buffer and left on ice for 10 minutes. Lysates were centrifuged (14,0000 rpm, 10 min at 4°C) and the detergent-soluble fraction-containing supernatant was collected. The pellets were resuspended in High Salt Buffer (HSB) (10mM Tris-HCl, 140mM NaCl, 1.5M KCl, 5mM EDTA, 0.5% Triton-X100 in ddH20) containing PPI and homogenized via glass douncer, followed by 1h incubation on a shaker at 4°C. Pellets (detergent-insoluble fraction) were centrifuged (14,0000 rpm, 20 min at 4°C), resuspended in PBS/EDTA (5mM EDTA in 1x PBS) and sonicated. The percent detergent-soluble K17 was calculated based on fluorescence of western blot bands using the formula: % sK17 = detergent-soluble/(detergent-soluble + detergent-insoluble).

### Cell proliferation assay

Cells were seeded into 96-well plate in triplicates. For each time point, 50 µl of 0.5% crystal violet (Feoktistova et al. 2016) was added and incubated for 20 minutes at RT. Excess crystal violet was removed by washing and the plate was air-dried. The remaining crystal violet was solubilized with 200 µl methanol and the optical density was read at 570 nm.

### Cell cycle analysis

Cells were harvested and stained with propidium iodide (Sigma-Aldrich) in Kreshan modified buffer. The percent cells at each cell cycle phase and apoptosis were measured by flow cytometry (BD Biosciences) and data were analyzed by ModFit LTTM (Verity Software House).

### Cell migration

Cell migration was assessed by wound-scratch assay. Cells were seeded followed by serum-starved in 1% FBS DMEM media. Scratch was performed using a 200 µl pipette tip, and wells were carefully washed with 1% FBS DMEM. Wound areas were imaged at subsequent time-points and measured using ImageJ software (NIH, Bethesda, MD, USA).

### Cell invasion

Cell invasion was assessed by Matrigel invasion chamber, using the Corning® BioCoat™ Matrigel^®^ Invasion Chamber (Corning, NY). Briefly, the inserts were rehydrated with serum-free DMEM for 2h. Cells previously serum-starved with 1% FBS DMEM media for 24h were trypsinized, washed in serum-free DMEM and added at a concentration of 1.5 x10^5^/ insert. Inserts were placed in a 24-well plate containing 10% FBS DMEM and incubated for 48h. Cells were fixed in 70% ethanol, air-dried, and then stained with 0.2% crystal violet. Rinsed and air-dried inserts were imaged at 10x and 5 fields were captured for each insert. Cell counts were averaged.

### RNA isolation, RT-PCR, and qRT-PCR

Total RNA was extracted with TRIzol reagent (Life Technologies), per manufacturer’s instructions. cDNA synthesis was performed using the Multiscribe method (ThermoFisher Scientific), using 1 µg of RNA as a template for cDNA generation. RT-PCR reaction was performed using custom TaqMan Array plates (ThermoFisher Scientific), pre-loaded with primers. TaqMan Universal Master Mix II, no UNG was used and qRT-PCR was programmed using an QuantStudio 3 real-time PCR system (ThermoFisher Scientific). Data were normalized by expression level of each sample as previously described (Schmittgen and Livak 2008).

### Computational analyses

K17 serine hotspot similarity between vertebrate species and human keratins was performed with MUSCLE v3.6 on the multiple sequence alignment NCBI ‘HomoloGene’ server, using option: ‘-maxiters 2’. K17 serine 10-13 site phosphorylation was predicted using NetPhos (Blom et al. 2004), GPS 5.0 (Wang et al. 2020) and Kinexus (Safaei et al. 2011) with default settings for all predictions. Data analyses and plotting were performed using R v4.0.3 or PRISM v8.2.1.

### Selection of genes enriched in the hepatic metastases of human PDAC samples

Microarray data from GEO accession 19280 were background-corrected, normalized, and summarized via the ‘rma’ function from the ‘oligo’ R package. Hepatic metastasis and primary pancreatic tumor samples were processed for subsequent differential expression analysis by applying a linear model to all genes to contrast metastatic and primary pancreatic tumor samples. Genes for which the median intensity < 4 (decided following visual inspection of median intensities) were excluded from differential expression analysis. The most significantly differentially over-expressed genes in hepatic metastases versus primary tumors, and 96 genes from the TaqMan^TM^ Array Human Tumor Metastasis plate (ThermoFisher Scientific) were evaluated for correlaiton with *KRT17* expression in primary pancreatic tumor sequencing datasets as reported in Moffitt *et al*, TCGA, and ICGC. The 20 genes with the strongest mean correlations with *KRT17* expression across the three datasets, and two genes (*CXCL10* and *DTYMK*) that we previously identified in cell lines to be associated with *KRT17* expression (unpublished observations) were chosen as potential hepatic metastasis-enriched genes to screen in our cell line model.

## Acknowledgments

The authors thank the Stony Brook University Cancer Registry (X. Barzilay) and Stony Brook Cancer Center Biorepository for assistance obtaining tissue specimens (Dr. R. Kew), Dr. Pierre Coulombe for the K17 antibody (University of Michigan), the Stony Brook University Flow Cytometry core facility (Todd Rueb and Rebecca Connor) and the Memorial Sloan Kettering Comprehensive Cancer Center Microchemistry and Proteomics core facility (Ronald C. Hendrickson). Dr. Ali Akalin provided human PDAC sections from the University of Massachusetts School of Medicine to support immunohistochemical studies.

## Finical support

The presented work was supported by: NCI K99-R00 CA226342-01 (L.F.E-H), William Raveis Charitable Fund / Rachleff Innovator of the Damon Runyon Cancer Research Foundation (L.F.E-H), AACR AACR Career Development Award to Further Diversity, Equity, and Inclusion in Pancreatic Cancer Research (L.F.E-H), the Stony Brook University Marvin Kuschner endowment (K.R.S), the Pancreatic Cancer Action Network Translational Research Grant (K.R.S), the National Pancreas Foundation Research Grant (C.V.L) and the National Institutes of Health Loan Repayment Program (C.V.L).

## Author contributions

Experiment conception/design: RRK, MTG, LRP, CVL, CHP, NTS, KRS, LFEH Methodology development: RRK, MTG, LRP, CVL, CHP, LTH, NTS, KRS, LFEH Data acquisition: RRK, MTG, LRP, CVL, CHP, LTH, TB, NTS, KRS Data analysis and interpretation: RRK, MTG, LRP, CVL, CHP, LTH, TB, NTS, KRS, LFEH, Writing of manuscript: RRK, MTG, KRS, LFEH Review of manuscript: All authors Study supervision: NTS, KRS, LFEH

## Supplementary figures

**Supplementary Figure S1.**
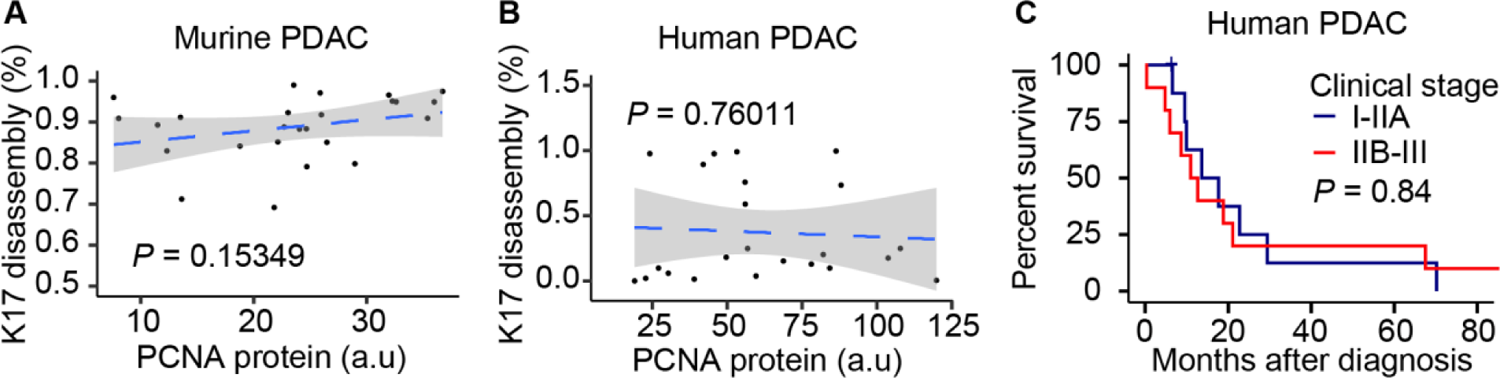
Detergent-soluble K17 is negatively prognostic in human and murine PDACs. **A-B.** Linear regression showing that K17 disassembly is independent of PCNA expression in mouse tumors *n*= 24 tumors (**A**) and human tumors *n*= 23 tumors (**B**). Significance determined by Spearman correlation test. **C.** Kaplan-Meier curves show the overall survival of human patients included in this study, according to clinical stage. Significance determined using Log-rank Mantel–Cox test.

**Supplementary Figure S2.**
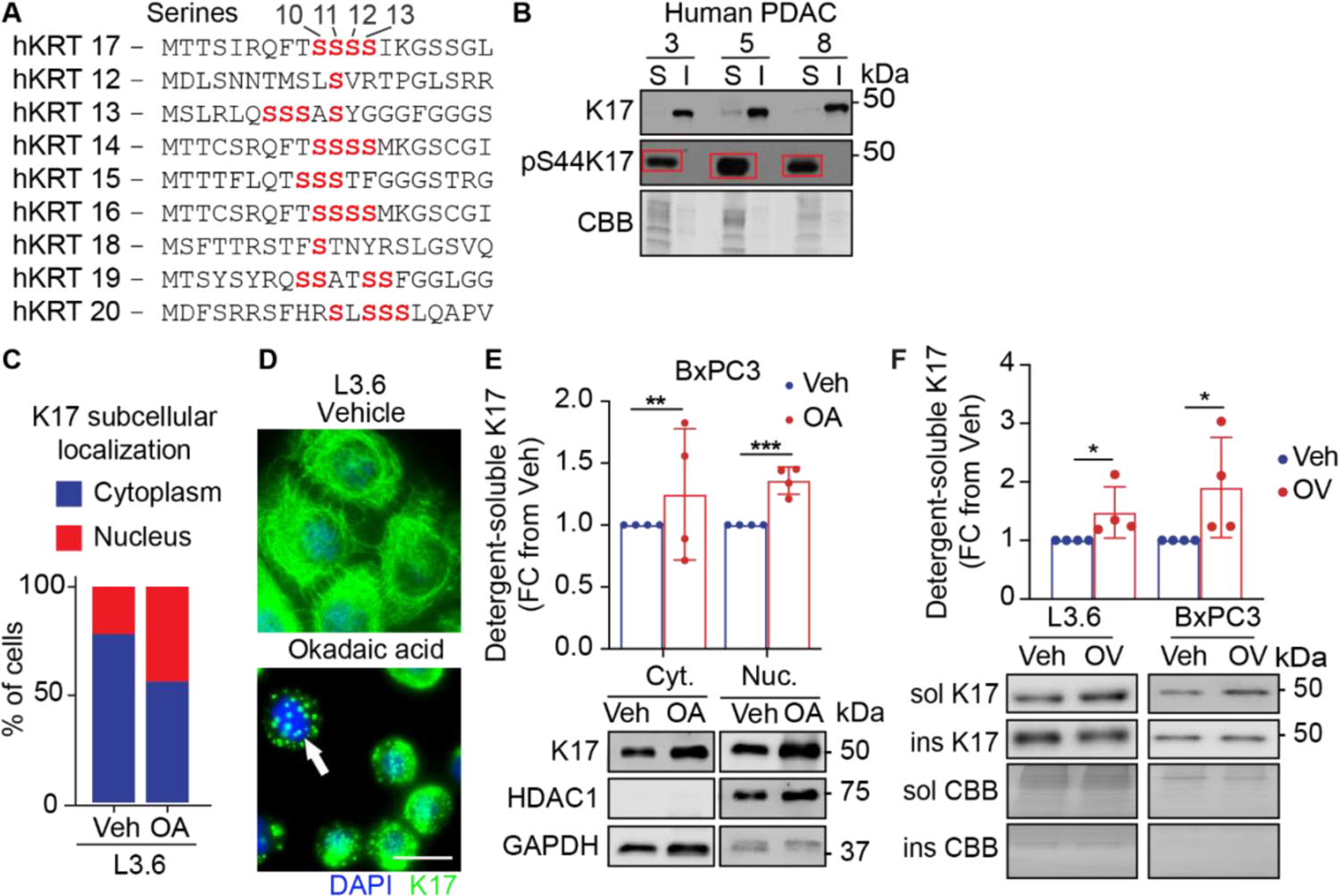
K17 is differentially phosphorylated in its detergent-soluble state, and N-terminal serine phosphorylation promotes disassembly. **A.** Alignment of type I human keratins polypeptide show similarity in the S10-13 hotspot. **B.** Serine 44 phosphorylation is highly enriched in detergent-soluble portions of the K17 protein in 3 representative human PDACs. **C-D.** Subcellular localization of K17 after okadaic acid (OA) treatment. Immunofluorescence stains (**C**) and quantifications (**D**) in L3.6 cells. Scale =5μm. **E**. Western blot depicting detergent-soluble and detergent-insoluble K17 after treatment with okadaic acid (OA) treatment. Treatment induces disassembly of K17 in L3.6 and BxPC3 cells. Data are mean ± *s.d, n* = 3 independent repetitions. Student’s *t*-test. **F.** Western blot depicting detergent-soluble and detergent-insoluble K17 after treatment with sodium orthovanadate (OV). Data are mean ± *s.d, n* = 3 independent repetitions. Student’s *t*-test. Coomassie Brilliant Blue (CBB), \**P*<0.05, \*\**P*<0.01, \*\*\**P*<0.001.

**Supplementary Figure S3.**
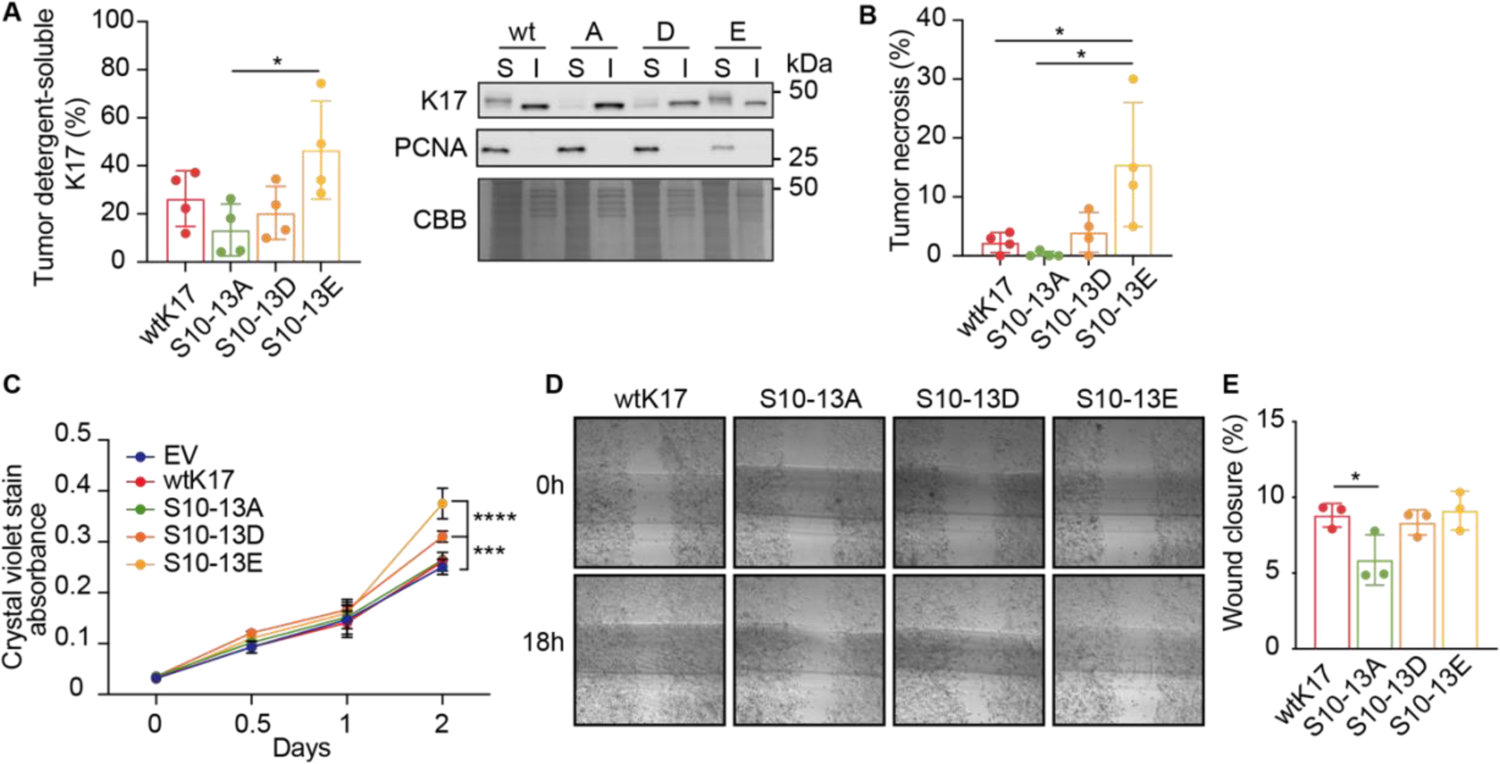
K17-phosphorylation impacts tumor growth and metastases *in vivo*. **A.** Percent detergent-soluble K17 in the tumors derived from GOPF S10-13E K17 wtK17 and LOPF S10-13A K17. Data are mean ± *s.d, n* = 4 independent PDACs. Student’s t-test. **B.** Percent tumor necrosis from PDACs derived from GOPF S10-13E K17 wtK17 and LOPF S10-13A K17. Data are mean ± *s.d, n* = 3 independent repetitions. Student’s t-test. **C.** Proliferation curve of K17-phosphorylation gain-of-function mutants (S10-13D and S10-13E), loss of function mutant (S10-13A) and wild type (WT) K17. **D-E.** Wound-closure assay (4x-**D**) and respective quantification (**E**) show that phosphorylation of K17 in S10-13 region enhances migration of PDAC cells. Coomassie Brilliant Blue (CBB), detergent-insoluble (ins, I), detergent-soluble (sol, S). \**P*<0.05, \*\*\**P*<0.001, \*\*\*\**P*<0.0001.

